# Freshwater Blackworms Solving a Maze

**DOI:** 10.64898/2026.10.13.759000

**Authors:** Andrew Adamatzky

**Affiliations:** Unconventional Computing Laboratory, University of the West of England, UK

**Keywords:** maze navigation, annelid behaviour, spatial statistics, behavioural mapping, exploratory locomotion

## Abstract

We investigated the behaviour of freshwater blackworms (*Lumbriculus variegatus*) in a circular maze containing food in its central chamber. Analysis of 11 time-lapse recordings showed that all tested worms entered the central chamber, although first-arrival times and spatial exploration patterns varied substantially. Registered occupancy maps revealed non-uniform use of corridors and chambers. Because the study did not include a no-food control, the observations demonstrate central-chamber entry and spatial exploration but do not by themselves establish food localisation or chemotaxis. The assay provides a reproducible framework for studying annelid exploration and for future controlled comparisons of sensory cues and maze complexity.

## 1. Introduction

The maze problem is one of the classical problems in mathematics, graph theory, and computer science. In its simplest form, the problem consists of finding a route between a designated source and destination in a geometrically constrained environment. To solve a maze is to find a route from the source site to the destination site. More generally, maze solving is closely related to shortest-path, path-planning, and navigation problems that arise in transportation systems, communication networks, robotics, logistics, and autonomous agents. Classical approaches rely on algorithmic procedures, such as graph-search algorithms, which operate on explicit representations of the environment and compute routes using symbolic manipulation of information.

Because maze solving is closely related to shortest-path computation, a variety of physical, chemical, and biological maze solvers have been developed. For example, a maze can be solved by representing its geometry by a network of electrical resistors [1] or a fluidic network [2]. A single human cell can solve a maze by crawling in a self-sustaining gradient generated through external and internal processes [3]. The slime mould *Physarum polycephalum* solves a maze by following gradients of attractants emitted by a food source placed in a central chamber [4]. Passive and active particles can solve mazes while relying on surface-tension-driven phenomena at the air–liquid interface [5]. Shortest-path computation in a maze of conductive channels can also be implemented using electrical current [6].

Freshwater blackworms (*Lumbriculus variegatus*) are increasingly used to investigate behaviour, physiology, pharmacology, environmental stress, and collective biophysics. Established assays quantify endpoints such as feeding, body reversal and stereotyped movement, dorsal-vessel pulsation, withdrawal or escape responses, and aggregation or disentanglement [7, 8, 9, 10, 11]. These assays are valuable for toxicology and pharmacology, but they do not test how an individual searches through a branched, geometrically constrained environment or how its occupancy is distributed across that environment. A maze assay therefore adds a spatially resolved behavioural endpoint that can be used to examine interactions among exploratory locomotion, boundary following, environmental cues, and geometry.

Our interest in maze-solving behaviour by invertebrates stems from earlier studies of medicinal leeches (*Hirudo verbana*), which demonstrated that organisms possessing relatively simple nervous systems can exhibit non-trivial exploratory behaviour in geometrically constrained environments [12, 13]. The present study extends this general experimental approach to a small, detritivorous aquatic annelid with different locomotor and sensory characteristics. The objective was not to infer a specific sensory mechanism, but to determine whether individual blackworms placed at the periphery would enter the central chamber and to quantify first-arrival time and spatial occupancy. This distinction is important because a central food pellet was present in all trials and no no-food control was performed.

## 2. Methods

Freshwater blackworms (*Lumbriculus variegatus*) were obtained from Blades Biological Ltd. Individual chronological age and fragmentation history could not be determined and worms were therefore not assigned an age. The archived experimental record does not contain sufficiently detailed pre-trial husbandry information to specify culture density, feeding regime, water temperature, pH, hardness, or dissolved oxygen; this limits exact replication and is acknowledged below.

PLA maze templates (Fig. 1a) were placed in sterile 120 × 120 mm square Petri dishes and submerged in municipal tap water supplied in Bristol (Fig. 1b). Bristol Water reports that its drinking water is treated and monitored under the Water Supply (Water Quality) Regulations 2016, that supplies across its service area are moderately hard to very hard because of dissolved calcium and magnesium, and that residual chlorine at customers’ taps is typically 0.1–0.8 mg L^−1^; chloramine is not added [14, 15, 16]. These supplier data provide general context only: no contemporaneous measurements of pH, hardness, conductivity, dissolved oxygen, or chlorine were made for the experimental water, and any pre-use conditioning was not documented. Figure 1a shows the exact two-dimensional maze geometry and can readily be used to reconstruct an STL model for replication. One worm was placed outside the template adjacent to its single entrance. The archived record establishes this common starting region but does not specify a fixed body orientation or distance from the entrance. A dog-food pellet (Arkwrights Complete with Beef Dry Dog Food, Gilbertson & Page Ltd) was placed in the central chamber in every trial; a no-food control was not included.

**Figure 1.**
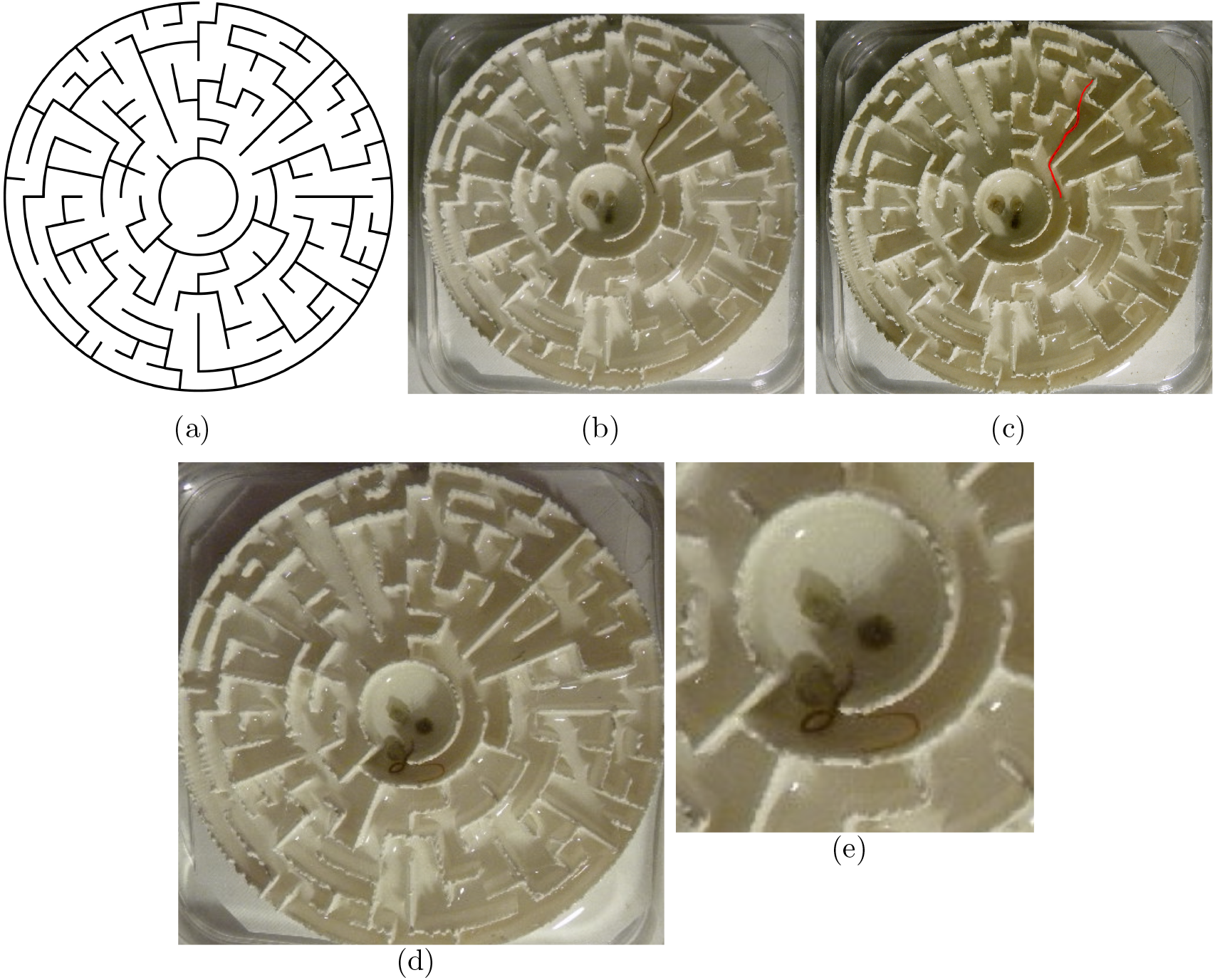
Experimental setup. (a) Geometry of the circular maze. (b) Experimental arena. (c) Example of automated worm detection (worm highlighted in red). (d) Worm in the central chamber; the image has been rotated to match the orientation in panels (a)–(c). (e) Enlarged region from panel (d).

Eleven independent trials were recorded with a FujiPix 6000 camera at one frame per minute. The sample size represented the complete set of usable recordings from this exploratory assay; no a priori power calculation was performed. Ambient laboratory illumination was used, but illuminance and spectral composition were not recorded. The underlying videos are archived on Zenodo [17].

Video processing and quantitative analysis were performed using custom software written in Processing, adapted from previous work [12, 13]. Scientific visualisations and statistical summaries were generated using Matplotlib, and registration and aggregation were implemented in custom Python routines. The analysis code is available from the corresponding author on reasonable request.

Each time-lapse recording was converted into a sequence of greyscale images. Because the maze remained stationary, a background image was estimated from the temporal median of sampled frames. Worm locations were identified as transient, dark filamentous structures relative to this background. Compact dark objects associated with the food pellet and static maze features were excluded, and only detections within the circular maze were retained. Small differences in maze placement were corrected by detecting the outer circular boundary and estimating its centre and radius; occupancy maps were then translated and scaled to a common coordinate system.

For each frame, worm pixels were assigned a value of one and other pixels a value of zero. Cumulative occupancy was calculated as

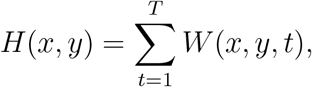

where *W* (*x, y, t*) denotes detected worm presence at pixel (*x, y*) and time *t*, and *T* is the number of frames. Registered maps were summed to describe common spatial patterns. For visual comparison, each displayed heatmap was normalised within its recording; the colour bar therefore represents relative cumulative occupancy from lower to higher values and not an absolute scale shared across recordings. Shannon entropy was calculated from each normalised occupancy distribution. Pearson’s correlation coefficient was used only as a descriptive measure of association between entropy and first-arrival time.

## 3. Results

Worms repeatedly moved between peripheral chambers and connecting corridors, and occupancy was non-uniform across the arena (Fig. 2). This pattern documents spatially heterogeneous exploration but does not distinguish attraction to food from thigmotaxis, undirected exploration, or other responses because food was present in every trial.

**Figure 2.**
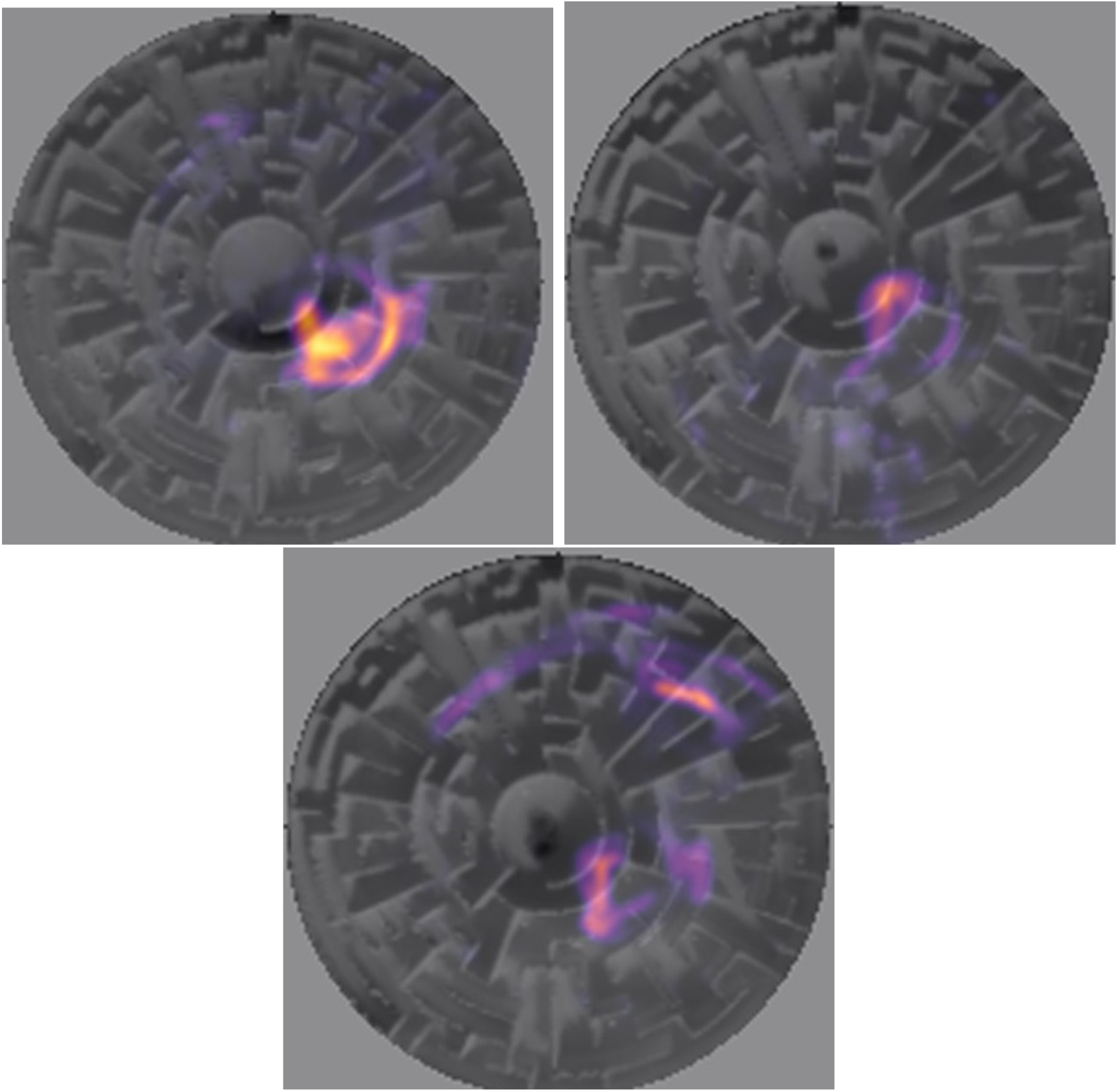
Examples of registered occupancy heatmaps obtained by integrating worm detections over all frames. Warmer colours denote higher cumulative occupancy after within-recording normalisation. The qualitative colour bar makes the direction of the scale explicit; colours should not be interpreted as a common absolute count across panels. Occupancy was spatially non-uniform.

The central chamber was entered in all 11 recordings (Fig. 1d–e). First-arrival time ranged from 90 to 1500 min, with a mean of 608.2 ± 443.8 min (mean ± SD) and a median of 480 min (Table 1). The value of 1500 min was the observed first-arrival time in that recording, not a predefined censoring limit. The large spread and small sample mean that these values should be treated as descriptive estimates rather than precise population parameters.

**Table 1:**
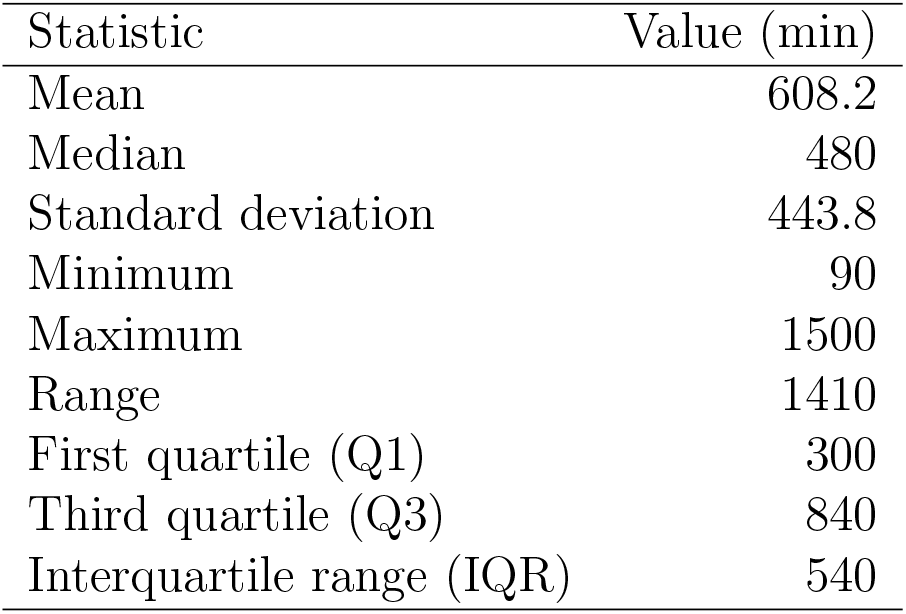
Descriptive statistics of first-arrival times to the central chamber (*n* = 11).

| Statistic | Value (min) |
| --- | --- |
| Mean | 608.2 |
| Median | 480 |
| Standard deviation | 443.8 |
| Minimum | 90 |
| Maximum | 1500 |
| Range | 1410 |
| First quartile (Q1) | 300 |
| Third quartile (Q3) | 840 |
| Interquartile range (IQR) | 540 |

Radial occupancy was greatest in peripheral and intermediate regions and decreased towards the centre (Fig. 3). The local peak immediately outside the central chamber may indicate prolonged passage through a geometrical bottleneck or hesitation near the chamber entrance; the present data cannot discriminate between these possibilities. Low central occupancy is also compatible with boundary-associated behaviour and negative phototaxis reported for blackworms, although illumination was not quantified in these experiments.

**Figure 3.**
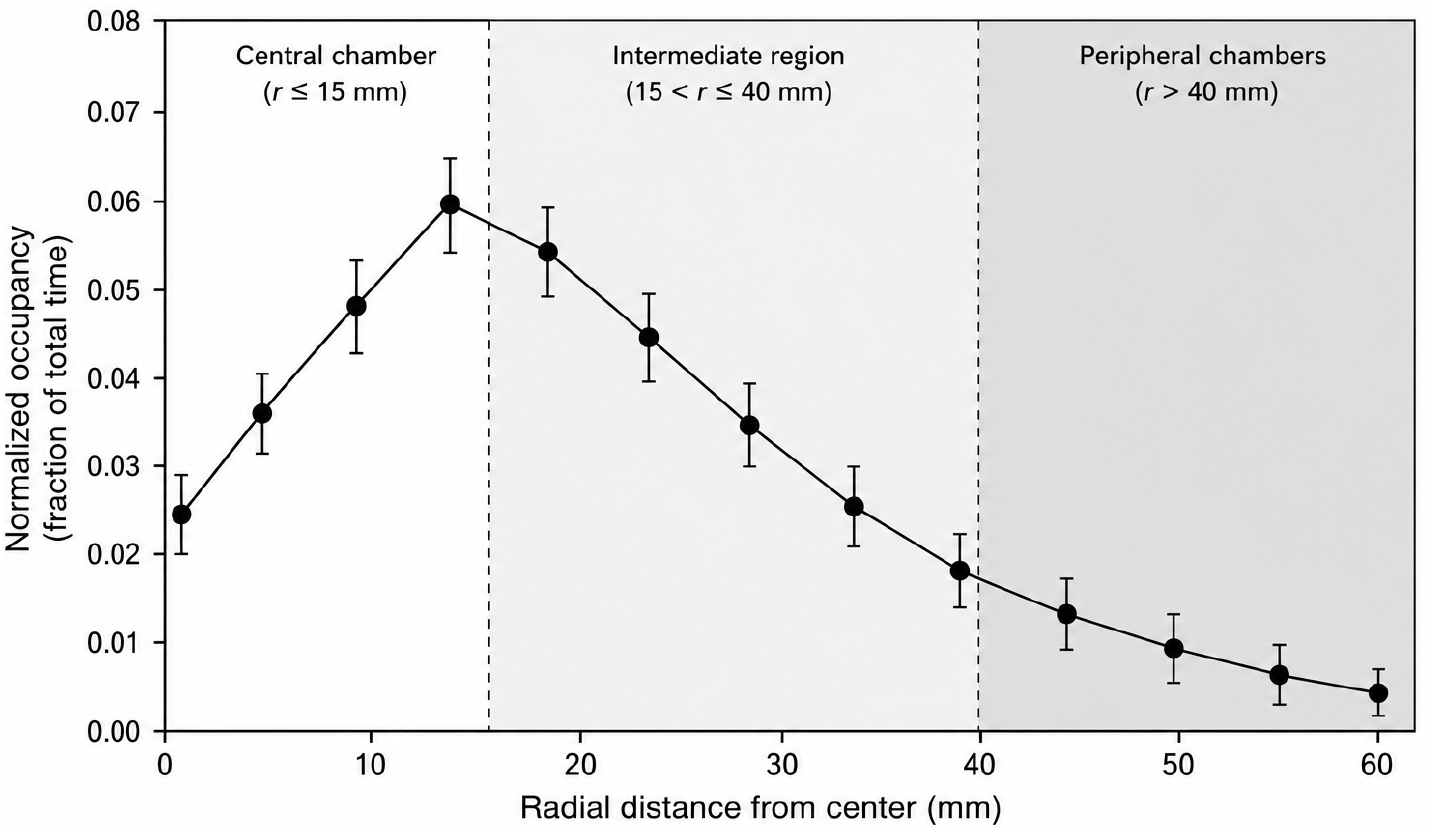
Radial occupancy profile of worm exploration in the circular maze. Occupancy was calculated from registered detections and expressed as the fraction of total observation time at each radial distance from the maze centre. Symbols show means across experiments and error bars show standard errors of the mean. Vertical dashed lines separate the central chamber, intermediate region, and peripheral chambers.

Entropy ranged from 11.54 to 14.15 bits, with a mean of 12.95 ± 0.75 bits. Higher values indicate occupancy distributed across more maze locations, whereas lower values indicate concentration within fewer corridors and chambers. The Pearson correlation between entropy and first-arrival time was *r* = 0.09, a negligible descriptive association in this sample (Fig. 4). It does not support a conclusion that route selection determined arrival time.

**Figure 4.**
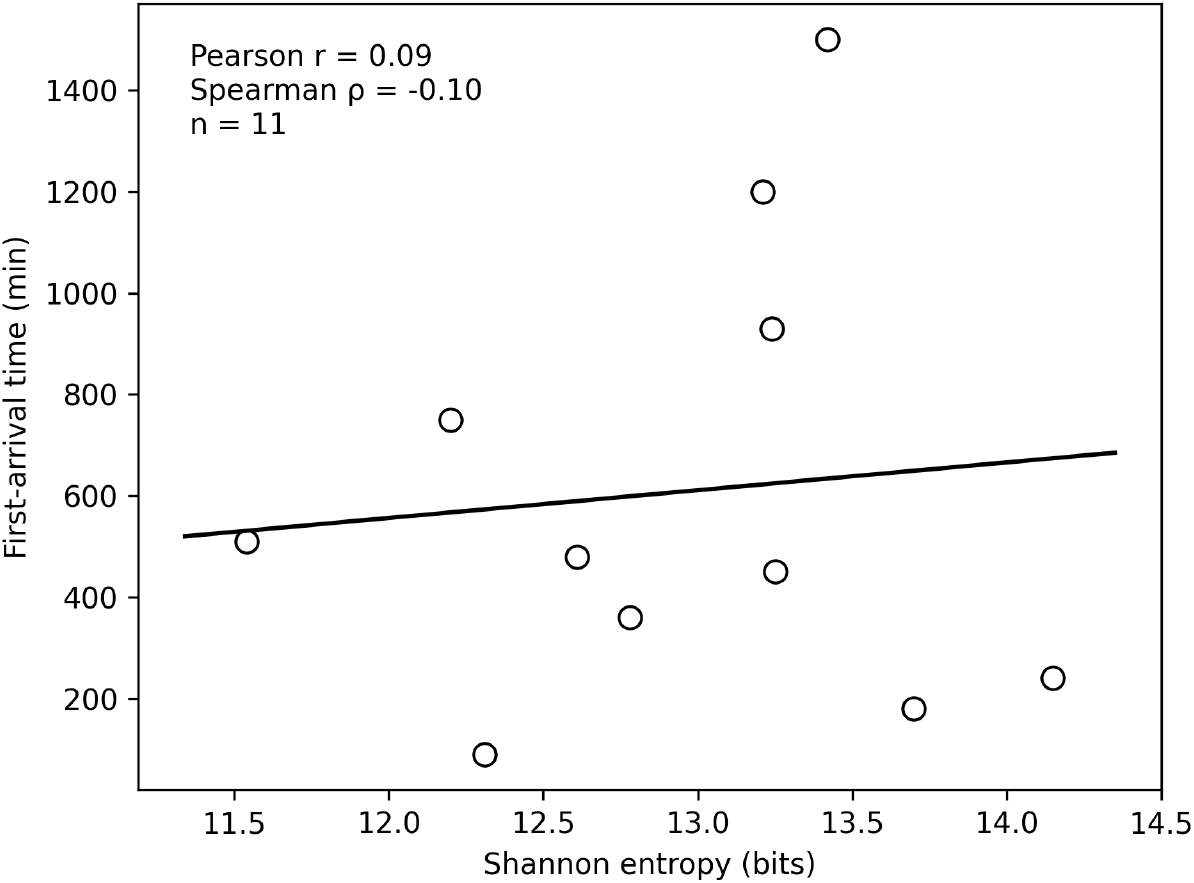
Shannon entropy versus first-arrival time. Each point represents one maze experiment.

## 4. Discussion

In this exploratory assay, every tested blackworm entered the central chamber of a circular maze, with substantial variation in first-arrival time and spatial occupancy. These observations establish that *L. variegatus* can traverse a branched maze from a peripheral starting region and provide quantitative endpoints for future behavioural experiments. They do not establish chemotaxis, food localisation, shortest-path selection, or goal-directed navigation. In particular, without an otherwise identical no-food condition, eventual entry may reflect undirected exploration, thigmotaxis, diffusion of food-associated cues, or a combination of these processes. Nutrient diffusion was not measured, so the presence, strength, and spatial form of any chemical gradient remain unknown.

The preference for peripheral and intermediate regions is compatible with the strong influence of boundaries on annelid locomotion and with the negative phototactic behaviour of blackworms. The peak immediately outside the central chamber may reflect a geometrical bottleneck or behavioural hesitation rather than attraction. Controlled illumination, tracking of approach and retreat events, and explicit measurements or manipulation of dissolved chemical gradients would be needed to resolve these alternatives. The negligible correlation between occupancy entropy and arrival time likewise indicates that these data do not identify a relationship between the overall spatial extent of exploration and arrival performance.

The assay complements established blackworm endpoints, including feeding, escape and withdrawal responses, stereotyped movement, dorsal-vessel pulsation, and collective aggregation [7, 8, 9, 10, 11]. Its potential contribution is a spatially structured test in which geometry, sensory cues, and behavioural state can be varied independently. This makes it relevant to behavioural ecology, environmental toxicology and pharmacology, as well as bio-inspired search and soft-robotics research. The comparison with prior leech studies is therefore methodological rather than a claim that the two taxa use the same sensory or decision-making mechanisms.

Several limitations should guide interpretation. The sample was small (*n* = 11), no a priori power analysis was conducted, and variability was high. There was no no-food control, no alternative maze geometry or complexity, and no measurement of nutrient diffusion. Although supplier information establishes that Bristol tap water is treated, monitored, mineral-rich, and chlorinated, the experimental water was not analysed contemporaneously and its conditioning was not documented; therefore, the precise exposure chemistry cannot be reconstructed. Pre-trial husbandry, precise starting orientation, and illumination were also not documented sufficiently for complete standardisation. These factors may affect blackworm physiology, phototaxis, wall following, and locomotion [9].

A confirmatory study should preregister its sample-size rationale and compare food-present and no-food conditions under defined water chemistry and lighting. It should standardise the start position, measure or manipulate putative chemical gradients, include mazes of different topology and complexity, and use sufficient replication to estimate effects reliably. These controls would permit direct tests of chemotaxis versus thigmotaxis and determine whether maze geometry changes arrival probability, latency, or route use.

## Data and software availability

The time-lapse recordings are available on Zenodo [17]. The exact maze geometry is shown in Fig. 1a and can readily be converted into an STL model. Custom analysis code is available from the corresponding author on reasonable request.

